# Microbial community-based production of single cell protein from soybean-processing wastewater of variable chemical composition

**DOI:** 10.1101/2022.08.02.502426

**Authors:** Ramanujam Srinivasan Vethathirri, Ezequiel Santillan, Sara Swa Thi, Hui Yi Hoon, Stefan Wuertz

**Author notes:** Correspondence to: Ezequiel Santillan and Stefan Wuertz.

## Abstract

The use of food-processing wastewaters to produce microbial biomass-derived single cell protein (SCP) is a sustainable way to meet the global food demand. Yet, despite the potential benefits of lower costs and greater resource recovery compared to pure cultures, bioconversion processes relying on microbial community-based approaches to SCP production have received scarce attention. Here, we evaluated SCP production from soybean-processing wastewaters under controlled reactor conditions using the existent microbial communities in these wastewaters. Six sequencing batch reactors of 4.5-L working volume were operated at 30 □ for 34 d in cycles consisting of 3-h anaerobic and 9-h aerobic phases. Four reactors received no microbial inoculum and the remaining two were amended with a 1.5 L of mixed culture from a prior microbial community-based SCP production. Microbial characterization was done via 16S rRNA gene metabarcoding. Influent wastewater batches had variable chemical characteristics but a similar microbial composition. Reactors produced more SCP when fed with wastewaters of higher soluble total Kjeldahl nitrogen (sTKN) content and a lower carbon-to-nitrogen ratio (sCOD:sTKN). The biomass protein yield ranged from 0.24 to 3.13 g protein/g sTKN, with a maximum protein content of 50%. An average of 92% of sCOD and 73% of sTN removal was achieved. Distinct microbial communities were enriched in all six bioreactors after 34 d, where the prevailing genera included *Azospirillum, Rhodobacter, Lactococcus, Novosphingobium*, and *Acidipropionibacterium*. In contrast, the microbial community of influent wastewaters was dominated by *Lactococcus* and *Weissella*. We showed that constituents in soybean wastewater can be converted to SCP through microbial community-based growth processes and demonstrated the effect of variable influent wastewater composition on SCP production.

## 1. Introduction

Microbial protein or single cell protein (SCP) consists of dried microorganisms with a high protein content along with fats, carbohydrates, vitamins, and minerals (Suman et al., 2015). It represents a promising alternative to fishmeal as a protein source in aquaculture, especially when raising sea bass, Atlantic salmon, rainbow trout, and whiteleg shrimp (Jones et al., 2020). Producing SCP from wastewaters would help alleviate the environmental impact caused by both traditional agricultural food production (Owsianiak et al., 2022) and wastewater treatment and disposal (Durkin et al., 2022; Spiller et al., 2020). SCP also has the potential to outperform staple crops in terms of protein yield per land area (Leger et al., 2021). According to estimates, by 2050, the adoption of microbial protein could reduce the required cropland area, global nitrogen losses from croplands, and agricultural greenhouse gas emissions worldwide by 6%, 8%, and 7%, respectively (Pikaar et al., 2018).

Microbial biomass may be produced by growing pure cultures (axenic SCP) or a mixture of strains or taxa (microbial community-based SCP), provided they can use wastewater as a source of carbon, energy and reducing power. A microbial community-based process has potential advantages over axenic culturing, such as a higher protein content due to synergistic interactions between different SCP-producing groups and utilization of various carbon and nitrogen sources in the substrate (Alloul et al., 2021a), process stability in terms of resistance and resilience to disturbances (Santillan et al., 2020b; Santillan et al., 2019a; Santillan & Wuertz, 2022), and the accumulation of intracellular components (Janarthanan et al., 2016). Further, this approach takes advantage of the microbial taxa already present in the wastewater by providing suitable growth conditions (Vethathirri et al., 2021). Indeed, the most suitable microbial organisms could be enriched within a mixed community to perform the targeted biotechnological process (Verstraete et al., 2022). Mixed microbial communities grown on wastewaters can meet the minimum dietary requirement of aquaculture animals (Vethathirri et al., 2021). However, information on the existent microbial communities in influent raw wastewaters and the enriched communities in such SCP production systems is scarce. Three types of bacterial consortia are used in microbial community-based systems, namely, aerobic heterotrophic bacteria, microalgae and aerobic heterotrophic bacteria, and purple phototrophic bacteria (Spiller et al., 2020; Vethathirri et al., 2021). Only the latter, which use light as energy source for the assimilation of organics and nutrients, have been directly enriched from wastewaters for SCP production (Hülsen et al., 2022). To our knowledge, the present study is the first to grow SCP based on aerobic heterotrophic bacteria present in the wastewater.

Compared to other wastewaters, food-processing wastewaters are of interest for microbial protein production due to their much lower content in pathogens, heavy metals, and other toxic contaminants. Chemically characterizing such wastewaters helps to assess their suitability for microbial biomass formation, with bioavailability of carbon- and nitrogen-containing organic compounds a key parameter (Vethathirri et al., 2021). In general, the chemical characteristics of food-processing wastewaters, including their carbon-to-nitrogen ratio (C:N), vary widely with production processes (Nayyar et al., 2021), making microbial community-based SCP production a challenging task. For it to be cost-effective and sustainable, the addition of external nutrients to maintain a suitable C:N should be avoided. Consequently, it is necessary to investigate how varying C:N affects both microbial growth and protein yield.

The overall aim of this study was to assess the production of microbial community-based SCP over time using different batches of the same source of soybean-processing wastewater. The objectives were to (1) microbially characterize the wastewater and community-based biomass; (2) determine suitable chemical characteristics in the feed to achieve a higher biomass yield and protein content; (3) characterize the amino acid content in the microbial protein produced; and (4) evaluate the effect of inoculation with a seed consortium on SCP production rate and yield. We showed that through microbial community-based growth, soybean-processing wastewaters can be converted directly into microbially derived protein that meets the amino acid requirements of aquaculture feed.

## 2. Materials and methods

### 2.1 Experimental design

Six 4.5-L bioreactors were operated as sequencing batch reactors on continuous 12-h cycles with intermittent aeration for 34 d, receiving wastewater from a soybean processing company in Singapore. Over the course of three months, nine 20-L carboys of wastewater from soybean soaking were collected at seven different occasions. Initially, two reactors were operated to test whether biomass could be enriched directly from soybean processing wastewater. Afterwards, a set of four reactors was operated to see if biomass could be again enriched directly from different batches of soybean processing wastewaters, and if having a starting inoculum would yield better SCP production than starting without it, receiving the same wastewater feed. Hence, four SBRs were operated without a starting inoculum: F_1-2_, F_3_, F_4-5_, and F_6-7_, whereas reactors IF_4-5_ and IF_6-7_ were started using 1.5-L inoculum from F_3_ after 43 days. Subscript numbers indicate the wastewater batch used as feed. Reactors IF_4-5_ and IF_6-7_ were comparable to reactors F_4-5_ and F_6-7_, respectively, which were fed with identical batches of wastewater but did not have a starting sludge inoculum.

### 2.2 Operational parameters and bioreactor arrangement

The reactor temperature was maintained at 30°C and sludge was continuously mixed at 375 rpm. Feeding phase occurred during the initial 5-10 min of a cycle, followed by alternating 180 min anoxic/anaerobic and 540 min aerobic phases. Cycles finished once soluble chemical oxygen demand (sCOD) of the mixed liquor was measured to be less than 400 mg/L, after which time the biomass was left to settle for 60 min and 1.85 L of supernatant was discarded. Thereafter, the reactor was filled with the same volume of soybean wastewater, starting a new cycle. This feeding scheme resulted in the following average hydraulic residence time (HRT) values for the six bioreactors operated: 9.7 d (F_1-2_), 14.6 d (F_3_), 8.4 d (F_4-5_), 8.4 d (IF_4-5_), 8.4 d (F_6-7_) and 8.4 d (IF_6-7_) (details in supplementary information). The pH ranged from 6.0 to 8.5 and the dissolved oxygen (DO) concentration was controlled between 0.2 and 0.5 mg/L during the aerobic phase. Each of the SBRs employed in this study was equipped with a magnetic stir plate to ensure mixed liquor homogeneity, a pair of EasySense pH and DO probes with their corresponding transmitters (Mettler Toledo), a dedicated air pump, a dedicated feed pump, a solenoid valve for supernatant discharge, and a surrounding water jacket connected to a re-circulating water heater. The different portions of the cycle were controlled by a computer software specifically designed for these reactors (VentureMerger, Singapore). Water chemical analyses were done as described in Santillan et al. (2021) (details in SI).

### 2.3 Biomass protein analysis

The protein content of the biomass was determined by quantifying amino acids using high-performance liquid chromatography (HPLC). First, 0.05 g of freeze-dried biomass was mixed with 5 mL of 6 M HCl and flushed with nitrogen gas for about 50 s. Then the samples was digested using heating block at 110°C for 22 h followed by filtration with a 0.22 µm membrane filter after cooling down. The solution was vacuum evaporated to dryness at 38 °C and 25 Pa in a rotary evaporator and re-dissolved in a volumetric flask with 5 mL of 0.1 M HCl. It was centrifuged for 40 min at 10,000 g and 4°C. The supernatant was filtered using a 0.22 µm membrane filter and the filtrate maintained at 4°C until derivatization (Chen et al., 2019). Pre-column derivatization was used with o-phthalaldehyde and 9-fluorenylmethoxycarbonyl (FMOC-Cl) via an autosampler (SIL -30 AC Autosampler). Seven and a half microliter of sample or standard was mixed with 45 µL of mercaptopropionic acid and 22 µL of OPA for 1 min in online derivatization. After mixing 8 µL of FMOC for 2 min, 5 µL of 0.1 M HCl was added. The reaction mixture was injected into an HPLC (Prominence UFLC, Shimadzu, Japan) equipped with a UV diode array detector (DAD, SPD-M20A), and detected at a wavelength of 338 nm for primary and 266 nm for secondary amino acids after passage through a Shimadzu Shim-pack Scepter C18 column (3µm, 3.0 × 150 mm). The flow rate was adjusted to 0.8 mL/min and the column temperature was set to 40°C. Gradient programs were applied for HPLC analysis.

### 2.4 Microbial analysis

Each soybean wastewater feed batch was subsampled once for microbial analysis and sludge samples were collected thrice a week from each reactor. A 50-mL feed sample was centrifuged at 10000 rpm for 3 min and 45 mL of supernatant discarded resulting in a tenfold increase in biomass concentration. Aliquots of 2 mL of concentrated wastewater and 2 mL of sludge samples were stored in cryogenic vials at −80°C for DNA extraction as previously described (Santillan et al., 2019b). Bacterial 16S rRNA amplicon sequencing was done in two steps as described in Santillan et al. (2020b). Primer set 341f/785r targeted the V3-V4 variable regions of the 16S rRNA gene (Thijs et al., 2017). The libraries were sequenced on an Illumina MiSeq platform (v.3) with 20% PhiX spike-in and at a read-length of 300 bp paired-end. Sequenced sample libraries were processed following the DADA2 bioinformatics pipeline (Callahan et al., 2016). DADA2 allows inference of exact amplicon sequence variants (ASVs) providing several benefits over traditional OTU clustering methods (Callahan et al., 2017). Illumina sequencing adaptors and PCR primers were trimmed prior to quality filtering. Sequences were truncated after 280 and 255 nucleotides for forward and reverse reads, respectively, length at which average quality dropped below a Phred score of 20. After truncation, reads with expected error rates higher than 3 and 5 for forward and reverse reads were removed. After filtering, error rate learning, ASV inference and denoising, reads were merged with a minimum overlap of 20 bp. Chimeric sequences (0.77% on average) were identified and removed. For a total of 81 samples, 47034 reads were kept on average per sample after processing, representing 52% of the average input reads. Taxonomy was assigned using the SILVA database (v.138) (Glockner et al., 2017). The adequacy of sequencing depth after reads processing was corroborated with rarefaction curves at the ASV level (Figure S1).

### 2.5 Fluorescence in situ hybridization

Microbial characterization was further supported by fluorescence in situ hybridization (FISH) using probes for the domain bacteria and selected core SCP taxa. Sludge samples were amended with 4% paraformaldehyde and placed on ice for 2-3 h. The fixed samples were washed with 1× phosphate-buffered saline (PBS) solution and stored in a mixture of 1×PBS and ethanol (1:1) at −20 °C until use. The cells were allowed to dry on microscopic slides and dehydrated in an ethanol series of 50, 80 and 96% for 3 min each. Hybridization buffer (0.9 M NaCl, 20 Mm Tris-HCl, 35% formamide, 0.01% sodium dodecyl sulfate) and probes were added to detect microorganisms of interest. Eubmix (Eub338, Eub 338ll, Eub 338lll) targets most bacteria (Daims et al., 1999) and AZOI 655 probe targets species belonging to *Azospirillum* (Stoffels et al., 2001). After hybridization, the slides were washed with warm buffer for 10 min (0.9 M NaCl, 20 Mm Tris-HCl, 5 mM EDTA, 0.01% sodium dodecyl sulfate) and rinsed thoroughly with cold water. FISH images were acquired using a LSM780 confocal laser scanning microscope. The Zen software was used for image processing and graphical analysis (Carl Zeiss, German).

## 3. Results and Discussion

### 3.1 Microbial composition of influent wastewaters

Microbial characterization of the influent food-processing wastewaters helps to identify microorganisms that could potentially drive the microbial community-based SCP production (Vethathirri et al., 2021). The different batches of soybean-processing wastewaters used in this study had a similar microbial community, which was dominated by the phylum *Firmicutes* at a relative abundance of 96 ± 6.7% (Figure 1a). At a finer taxonomic level, *Lactococcus* and *Weissella* were the most abundant genera at relative abundances of 45 ± 16.6% and 36 ± 16.6%, respectively (Figure 1b). Strains of *Lactococcus* are used in cheese making and can produce vitamins such as B2 and K2 (Song et al., 2017), and *Weissella* has shown potential as a probiotic in food and pharmaceutical industries (Teixeira et al., 2021). In addition, *Leuconostoc* was identified in all soybean wastewaters at a relative abundance of 4.1–15.1% except for the sixth wastewater batch, which at 14 mg/L also had the lowest average sTKN content. The most abundant ASV at the species level was *Leuconostoc palmae* (up to 6.5% relative abundance). Several strains of *Leuconostoc* are of economic value due to their widespread application in dairy technology (Hemme & Foucaud-Scheunemann, 2004). Besides these three genera, *Streptococcus* was also found in all wastewater batches at a relative abundance of up to 16%. *Streptococcus thermophilus* is important in dairy industries that produce milk, cheese, and yogurt (Mora et al., 2002). *Lactobacillus, Enterobacter, Aeromonas*, and *Acinetobacter* occurred less frequently in influent wastewaters at a relative abundance greater than 1% included (Figure 1b). *Lactobacillus* can be used as a probiotic to cure several ailments, especially chronic liver diseases (Jeong et al., 2022), and strains of *Lactobacillus rhamnosus, Lactobacillus casei* and *Lactobacillus plantarum* may be explored as a source of natural antioxidants (Shori et al., 2022). Certain species of free-living *Enterobacter*, specifically *Enterobacter cloacae*, were found to be involved in symbiotic nitrogen fixation in plants such as wheat (Ji et al., 2020). Studying the microbial communities of influent wastewaters can also help spot potentially pathogenic *Aeromonas* strains associated with disease in humans and aquatic animals (Fernandez-Bravo & Figueras, 2020). In conclusion, the microbial characterization of influent wastewaters revealed low variability across batches and identified several taxa suitable for the potential production of both SCP and other valuable compounds.

**Figure 1.**
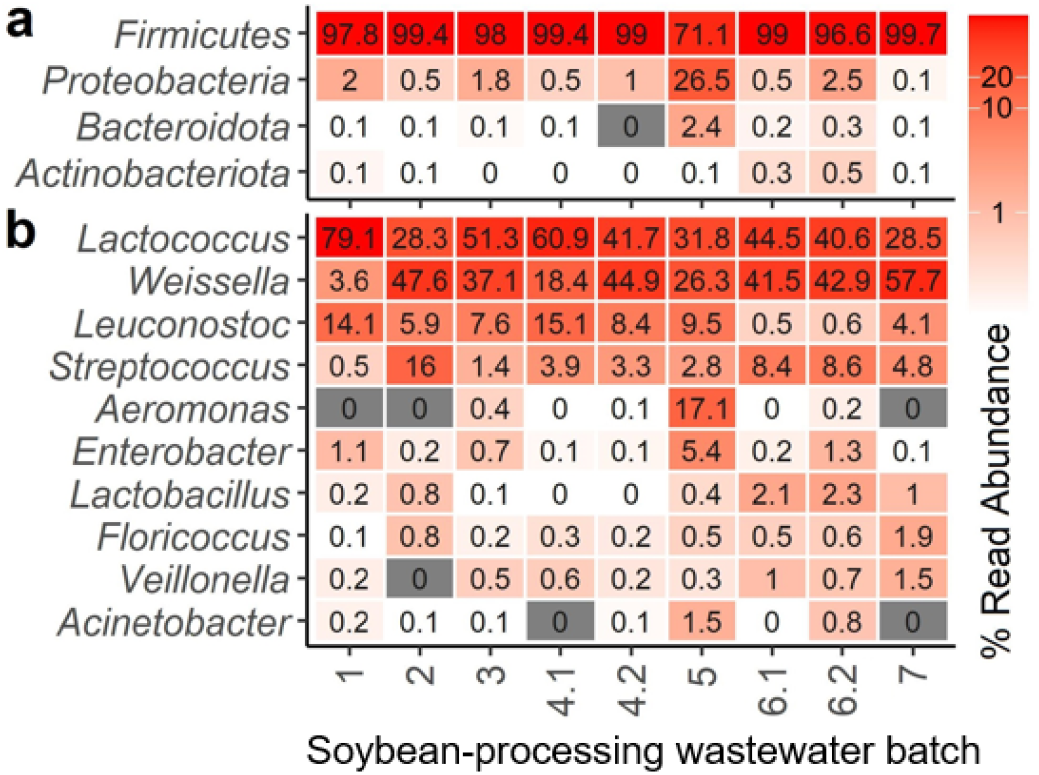
Microbial characterization of influent soybean processing wastewaters used for microbial community-based SCP production as assessed through 16S rRNA gene amplicon sequencing (n = 9). **(a)** Four most abundant phyla and **(b)** ten most abundant genera of nine wastewater batches were collected at seven different time points over three months.

### 3.2 Wastewaters with a lower sCOD:sTKN (C:N) and higher sTKN produced more SCP

While the microbial communities of different batches of soybean-processing wastewaters were similar at the phylum level, the chemical composition was variable, particularly the nitrogen content. The sTKN and sCOD:sTKN values were in the range of 13 −110 mg/L and 74 - 187, respectively, for the seven batches of wastewaters collected at different time points from the same soybean processing source (Table 1). According to the wastewater provider, this variability could be due to changes in the quantity of soybeans soaked in water for washing and the source of the soybeans being processed. In general, the darker the color of soybean-processing wastewater, the higher the sTKN level and the lower the sCOD:sTKN value, increasing its potential for SCP production (Vethathirri et al., 2021). To account for the variable N content in different batches of wastewater used in this study, we characterized them as either low (L) for sTKN = 13 - 61 mg/L or high (H) for sTKN = 62 - 110 mg/L) (see Tables 1 and 2).

**Table 1.** Chemical characteristics of soybean processing wastewaters used for microbial community-based SCP production.

| Soybean wastewater batch<sup>a</sup><br>(quality)<sup>b</sup> | sCOD<br>[mg/L] | sTKN<sup>c</sup><br>[mg/L] | sPO<sub>4</sub><sup>3-</sup>-P<br>[mg/L] | NH<sub>4</sub><sup>+</sup>-N<br>[mg/L] | NO<sub>2</sub><sup>-</sup>-N<br>[mg/L] | NO<sub>3</sub><sup>-</sup>-N<br>[mg/L] | TA<sup>d</sup><br>[mg/L] | TSS<br>[mg/L] | VSS<br>[mg/L] | pH | C: N<br>(sCOD:<br>sTKN)<sup>e</sup> | N:P (sTKN:<br>sPO<sub>4</sub><sup>3-</sup>-P)<sup>e</sup> |
| --- | --- | --- | --- | --- | --- | --- | --- | --- | --- | --- | --- | --- |
| 1 (H) | 9038 | 110 | 67 | 5.9 | 0.08 | 13.65 | 453 | 90 | 90 | 5.90 | 82.5 | 1.6 |
| 2 (L) | 2334 | 31 | 17 | 3.1 | 0.04 | 1.88 | 57 | 140 | 140 | 4.80 | 74.3 | 1.9 |
| 3 (H) | 8945 | 110 | 65 | 9.0 | 0.32 | 8.4 | 208 | 530 | 480 | 4.80 | 81.3 | 1.7 |
| 4.1 (L) | 5670 | 50 | 21 | 6.42 | 0.01 | 0.00 | 206 | 210 | 210 | 5.39 | 112.0 | 2.4 |
| 4.2 (L) | 4180 | 28 | 18 | 4.07 | 0.00 | 0.09 | 138 | 310 | 310 | 5.28 | 147.0 | 1.6 |
| 5 (L) | 3474 | 34 | 14 | 3.40 | 0.00 | 0.02 | 118 | 160 | 160 | 5.04 | 101.8 | 2.5 |
| 6.1 (L) | 2494 | 13 | 8 | 1.28 | 0.00 | 0.37 | 39 | 150 | 150 | 4.90 | 186.9 | 1.7 |
| 6.2 (L) | 2501 | 15 | 8 | 1.11 | 0.01 | 0.31 | 34 | 180 | 180 | 4.85 | 166.2 | 1.9 |
| 7 (H) | 7663 | 80 | 29 | 5.26 | 0.00 | 0.07 | 254 | 330 | 330 | 4.70 | 96.1 | 2.8 |

<sup>a</sup> A total of nine 20-L carboys were collected at seven different time points over three months.
<sup>b</sup> High quality (H) for sTKN = 62 - 110 mg/L and low quality (L) for sTKN = 13 - 61 mg/L.
<sup>c</sup> sTKN was calculated using the value of sTN, nitrate and nitrite: sTKN = sTN – nitrate - nitrite.
<sup>d</sup> Total alkalinity expressed as calcium carbonate
<sup>e</sup> Ratio of carbon and nitrogen in wastewater as calculated from chemical data.

**Table 2.** Microbial community-based SCP production through bioconversion of soybean processing wastewaters.

| Reactor | Day | Waste water batch (quality)<sup>a</sup> | sTKN [mg/L] | C: N [sCOD: sTKN] | Biomass production rate [g TSS/L<sub>R</sub>/d]<sup>b</sup> | Biomass yield_sCOD [g TSS/g sCOD] | Biomass yield_sTKN [g TSS/g sTKN] | Protein production rate [g SCP/L<sub>R</sub>/d] | Protein yield_sCOD [g SCP/g sCOD] | Protein yield_sTKN [g SCP/g sTKN] | R<sub>sCOD</sub><sup>c</sup> [%] | R<sub>sTN</sub><sup>c</sup> [%] |
| --- | --- | --- | --- | --- | --- | --- | --- | --- | --- | --- | --- | --- |
| F<sub>1-2</sub> | 1 – 23 | 1 (H) | 110 | 82.5 | 0.005 | 0.006 | 0.49 | 0.006 | 0.007 | 0.53 | 87 | 74 |
|  | 24 – 34 | 2 (L) | 31 | 74.3 |  |  |  |  |  |  |  |  |
| F<sub>3</sub> | 1 – 34 | 3 (H) | 110 | 81.3 | 0.042 | 0.053 | 4.28 | 0.010 | 0.012 | 1.00 | 90 | 68 |
| F<sub>4-5</sub> | 1 – 14 | 4.1 (L) | 50 | 112.0 | 0.013 | 0.021 | 2.55 | 0.004 | 0.007 | 0.82 | 95 | 75 |
|  | 15 – 29 | 4.2 (L) | 28 | 147.0 |  |  |  |  |  |  |  |  |
|  | 30 – 34 | 5 (L) | 34 | 101.8 |  |  |  |  |  |  |  |  |
| IF<sub>4-5</sub><sup>d</sup> | 1 – 14 | 4.1 (L) | 50 | 112.0 | 0.021 | 0.036 | 4.44 | 0.006 | 0.011 | 1.37 | 93 | 74 |
|  | 15 – 29 | 4.2 (L) | 28 | 147.0 |  |  |  |  |  |  |  |  |
|  | 30 – 34 | 5 (L) | 34 | 101.8 |  |  |  |  |  |  |  |  |
| F<sub>6-7</sub> | 1 – 14 | 6.1 (L) | 13 | 186.9 | 0.039 | 0.083 | 11.06 | 0.011 | 0.024 | 3.13 | 93 | 59 |
|  | 15 – 29 | 6.2 (L) | 15 | 166.2 |  |  |  |  |  |  |  |  |
|  | 30 – 34 | 7 (H) | 80 | 96.1 |  |  |  |  |  |  |  |  |
| IF<sub>6-7</sub><sup>d</sup> | 1 – 14 | 6.1 (L) | 13 | 186.9 | 0.040 | 0.089 | 11.64 | 0.009 | 0.021 | 2.76 | 94 | 88 |
|  | 15 – 29 | 6.2 (L) | 15 | 166.2 |  |  |  |  |  |  |  |  |
|  | 30 – 34 | 7 (H) | 80 | 96.1 |  |  |  |  |  |  |  |  |

<sup>a</sup> High quality (H) for sTKN = 62 - 110 mg/L and low quality (L) for sTKN = 13 - 61 mg/L.
<sup>b</sup> L<sub>R</sub> = reactor volume in liters. All reactors had a working volume of 4.5 L.
<sup>c</sup> Average nutrient removal efficiency calculated for each reactor based on feed and effluent characteristics (Table S1).
<sup>d</sup> Reactors IF<sub>4-5</sub> and IF<sub>6-7</sub> received a 1.5-L inoculum from reactor F<sub>3</sub>, while reactors F<sub>1-2</sub>, F<sub>3</sub>, F<sub>4-5</sub> and F<sub>6-7</sub> were started without inoculum.

The biomass in the six community-based SCP reactors was tracked as total suspended solids (TSS) concentration for 34 days (Figure 2a). The initial average biomass concentration of the four reactors that started without inoculum was the same as the average TSS of the soybean wastewater batches (233 mg/L), whereas the other two reactors that received inoculum had an initial average TSS of 575 mg/L. Despite having started without inoculum, reactor F_3_ showed a higher TSS concentration of 1967 mg/L on d34 than its offspring reactors IF_4-5_ and IF_6-7_, which started with an inoculum produced from F_3_. We attribute this difference to the favorable chemical characteristics of the wastewater batch used for F_3,_ which had high sTKN levels and a low sCOD:sTKN compared to other feeds (Table 2). On the other hand, the lowest TSS value on d34 was 227 mg/L, observed in reactor F_1-2_, which was initiated without inoculum. Further, this was also the only reactor that showed a declining trend in cell growth in the second half of the experiment (Figure 2a). This was due to the low sTKN level in the wastewater batch (31 mg/L) that was used after d24, which then affected the overall biomass production rate and yield (Table 2) of reactor F_1-2_. Although this batch had the lowest sCOD:sTKN of all the batches (Table 2), it did not support cell growth with the feeding regime used in this study. Hence, both a lower sCOD:sTKN and a sufficiently high sTKN concentration are needed to sustain community-based microbial biomass production.

**Figure 2.**
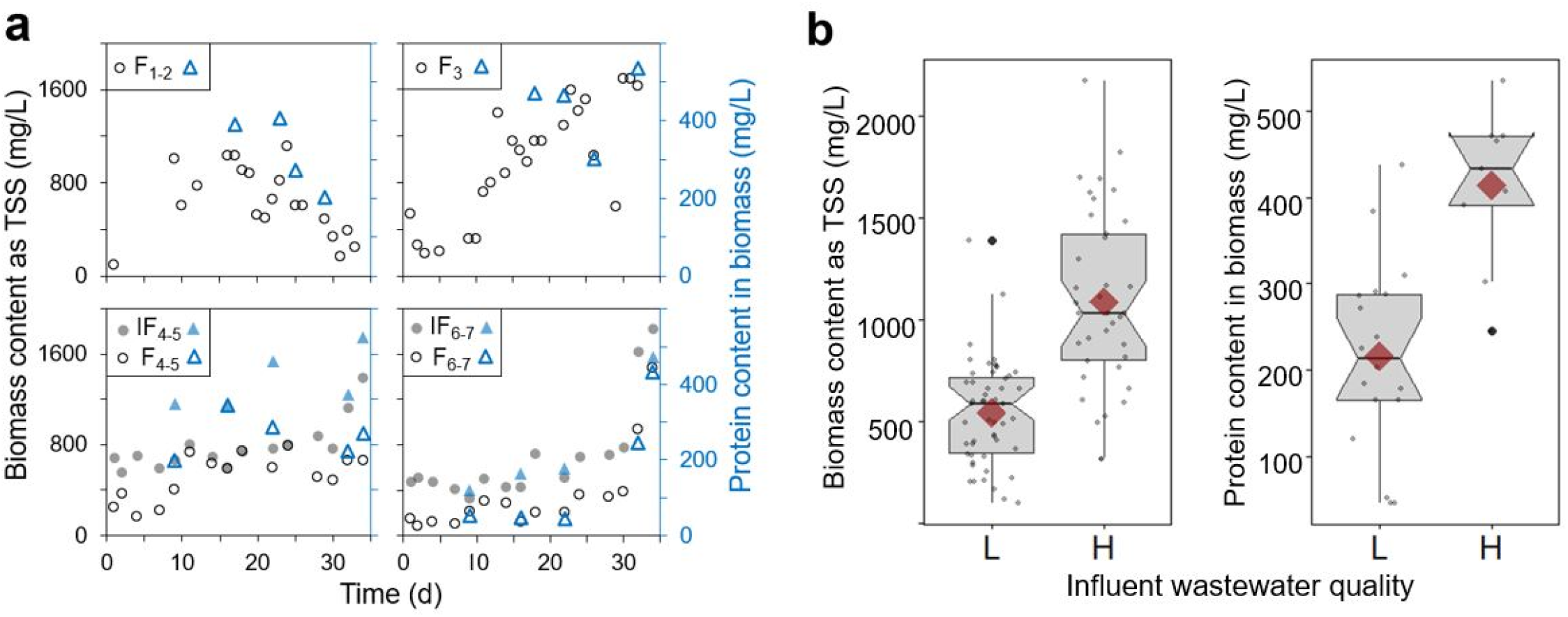
Biomass growth, biomass protein content and influent wastewater quality impact in six SCP-enriched bioreactors. **(a)** Biomass measured as total suspended solids (TSS, black circles), and protein content in the biomass via amino acid quantification using HPLC (blue triangles). Four reactors (F_1-2_, F_3_, F_4-5_, and F_6-7_) were initiated without inoculum and two (IF_4-5_ and IF_6-7_) received a 1.5-L inoculum from reactor F_3_ (subscript numbers indicate the wastewater batch used as feed). **(b)** Box plots of biomass content and protein content in biomass for low (L, sTKN = 13 - 61 mg/L) and high (H, sTKN = 62 - 110 mg/L) quality feed. Only data from day 5 onwards were included to exclude low TSS values during the reactor start-up period. Red diamonds display mean values and notches show the 95% confidence interval for the median; when notches do not overlap the medians can be judged to differ significantly (two-tailed Welch’s t-tests, p < 0.0001).

Amino acid analysis by HPLC was used to accurately determine protein levels in the dry microbial biomass (FAO, 2003; Vethathirri et al., 2021). The amount of protein in the biomass followed the pattern of TSS production (Figure 2a), with increasing or decreasing trends depending on the wastewater batch used in each bioreactor at a particular time of the study. An increase in biomass protein content was observed in reactor F_1-2_ from d1 to d23, in F_3_ from d1 to d34, in F_6-7_ from d22 to d34, and in IF_6-7_ from d22 to d34 due to the usage of feed wastewaters with a lower sCOD:sTKN and higher sTKN content (62 - 110 mg/L) (Figure 2a). A similar pattern was observed with other types of mixed-culture SCP biomass produced from photosynthetic bacteria using artificial sugar wastewaters of lower COD:TN (5 to 50) (Cao et al., 2021). On the other hand, biomass protein content was decreasing or stayed constant in reactors F_1-2_ (d24 to d34), F_4-5_ (d1 to d34), IF_4-5_ (d1 to d34), F_6-7_ (d1 to d21), and IF_6-7_ (d1 to d21) due to the usage of wastewaters with a higher sCOD:sTKN and lower sTKN content (13 - 61 mg/L) (Figure 2a) in our study. Hence, when reactors received wastewater with a lower sCOD:sTKN and a higher sTKN content, they produced significantly more biomass and protein (two-tailed Welch’s t-tests, p < 0.0001) than when they were fed with wastewater that had a high sCOD:sTKN and low sTKN content (Figure 2b).

The protein content in the microbial biomass is important for fish feed applications (Azim et al., 2008; Webster & Lim, 2002). Higher protein contents of 49.9, 49.5 and 48.7% as dry biomass were observed in reactors IF_4-5_, F_1-2_, and F_4-5_, respectively (Figure S2), regardless of the presence or absence of an initial inoculum. Similar protein percentages were observed in aerobic heterotrophic biomass produced from different food processing wastewaters (Muys et al., 2020). However, they were lower than reported values for other types of SCP biomass. For instance, purple non-sulfur bacteria (PNSB) growing on volatile fatty acids had a maximum protein content of 61% as dry cell weight (Peng et al., 2022), although this was measured with the modified Lowry method. Colorimetric protein assays, like the modified Lowry and Biuret methods, are not recommended for assessing the protein content in SCP production studies (Vethathirri et al., 2021) because they are susceptible to interferences that often lead to erroneous readings (Le et al., 2016). Lower protein contents of 29.3% and 29.1% as dry biomass were observed in reactors F_6-7_ and IF_6-7_, respectively, (Figure S2) towards the end of the study due to the usage of wastewater with a higher sCOD:sTKN and lower sTKN content until d29, despite the shift to a wastewater feed with a higher sTKN content afterwards (Table 2). Additionally, none of the reactors had a constant protein fraction during the bioconversion period, including reactor F_3_ which was fed with the same wastewater batch throughout the study. Our data showed that the highest protein fraction (∼ 50%) was achieved during 15 to 25 days.

### 3.3 Microbial community-based protein meets amino acid requirement for aquaculture feed

The SCP produced using several batches of soybean wastewaters either completely or partially met the demand of essential amino acids (EAA) in the aquaculture diet of trout (Ogino, 1980) and shrimp (Tacon et al., 2002) (Figure 3). Ten amino acids are regarded essential, because they are not produced by these aquaculture animals and thus need to be supplemented as part of their diet (Millamena, 2002). Microbial biomass produced in the six bioreactors had an EAA content of 38.8 ± 2.0% in protein and featured all EAAs, except for tryptophan which was not measured. Among them, leucine was found to be the most enriched in all bioreactors (9.2 ± 0.6% of protein). Leucine is a valuable branched chain amino acid supporting the growth, maintenance, metabolic, and physiological needs of fish (Ahmad et al., 2021). The dry biomass produced using seven distinct batches of soybean wastewaters was also rich in aspartic acid, glutamic acid, and alanine (Figure S3 and Figure S4). The same amino acids were also dominant in the existent microbial community (measured as dry cell mass) in soybean-processing wastewater (Figure S5). SCP produced from industrial wastewaters was reported to be suitable for replacing, at least partially, conventional fish meal (Hülsen et al., 2018b; Muys et al., 2020). When SCP produced from purple bacteria was used as shrimp feed, the animals exhibited higher tolerance of *Vibrio* pathogens and resistance to ammonia stress compared to a control group fed on conventional fish meal (Alloul et al., 2021b). Further, palatability of the microbial community-based SCP can be promoted by adding attractants or feed stimulants like betaine and free amino acids (Ajiboye et al., 2012) or by using feed additives like onion powder (Anwer et al., 2018). Hence, SCP produced by the growth of microbial communities can potentially be used as a protein source in the feed of aquaculture animals.

**Figure 3.**
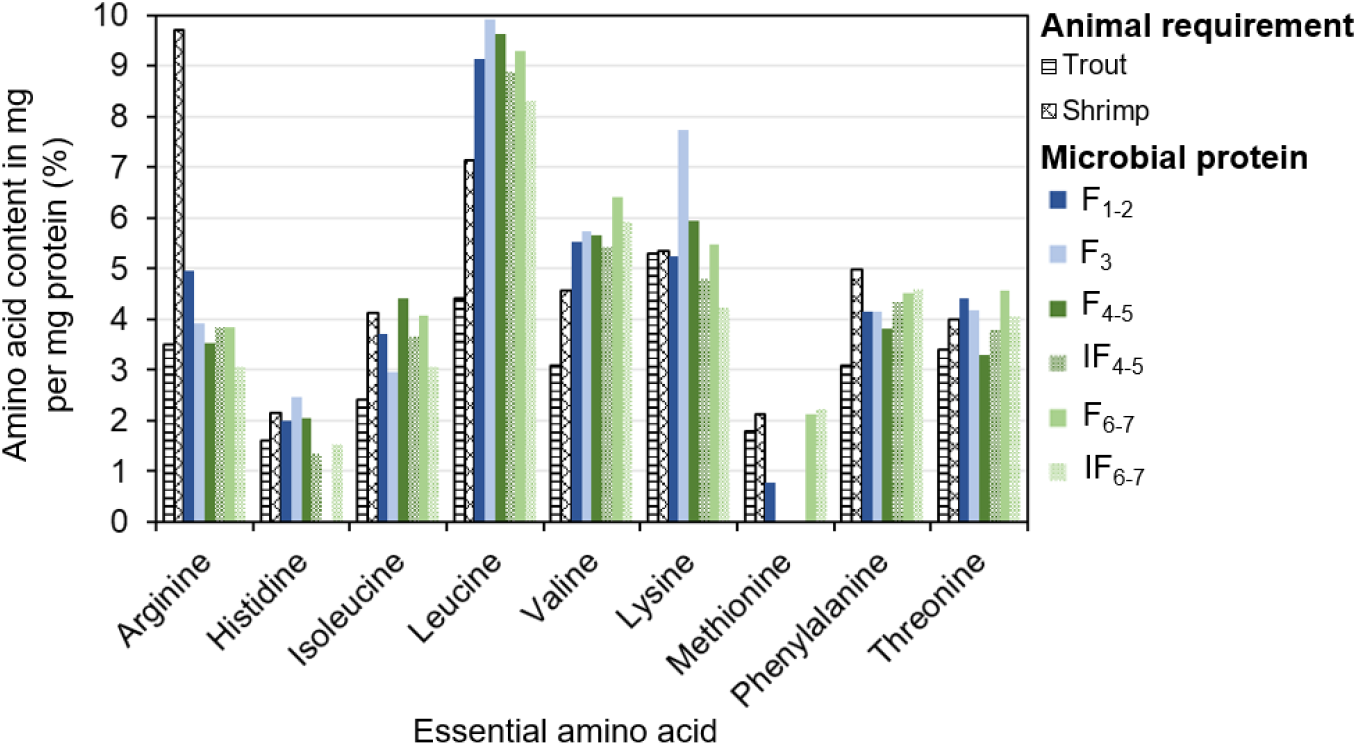
Essential amino acids in microbial community-based single cell protein produced from soybean processing wastewaters (solid/pattern-filled blue/green bars) and compared to the minimum requirements for trout and shrimp feed (pattern-filled black bars). Data available in Ogino (1980) and Tacon et al. (2002) were used to plot the minimum essential amino acid requirements of trout and shrimp, respectively. Four reactors were initiated without inoculum: F_1-2_ (solid-filled dark blue bar), F_3_ (solid-filled light blue bar), F_4-5_ (solid-filled dark green bar), and F_6-7_ (solid-filled light green bar). Two reactors received a 1.5-L inoculum from reactor F_3_: IF_4-5_ (pattern-filled dark green bar) and IF_6-7_ (pattern-filled light green bar). Subscript numbers indicate the wastewater batch used as feed. Data used for F_1-2_ and F_3_ correspond to samples collected on d29 and d32, respectively, while data used for the remaining reactors correspond to d34.

### 3.4 Impact of inoculum on microbial community-based SCP production

Microbial community-based synthesis of single cell protein from wastewaters has generally involved inoculation with a seed culture derived from previous experiments (Hülsen et al., 2018a; Hülsen et al., 2018b; Hülsen et al., 2020). In the present study, two out of six bioreactors received 1.5 L of a mixed microbial community that grew in a prior SCP experiment using the same source of soybean-processing wastewater. sCOD removal efficiencies (R_sCOD_) in these reactors, IF_4-5_ (R_sCOD_ = 93%) and IF_6-7_ (R_sCOD_ = 93%), were like those in their corresponding reactors, F_4-5_ (R_sCOD_ = 94%) and F_6-7_ (R_sCOD_ = 93%), respectively, which were operated without inoculum but received the same batches of soybean processing wastewater (Table 2). In contrast, the sTN removal efficiency (R_sTN_) of only one reactor, IF_4-5_ (R_sTN_ = 74%), was comparable to that of its counterpart reactor, F_4-5_ (R_sTN_ = 75%), whereas the sTN removal efficiency (R_sTN_) of the other reactor, IF_6-7_ (R_sTN_ = 88%) was higher than that in its counterpart reactor, F_6-7_ (R_sTN_ = 59%). We attribute this difference to the high-quality wastewater (sTKN = 80 mg/L) used to feed reactors F_6-7_ and IF_6-7_ from d29 to d34. Hence, inoculated reactors exhibited a higher nitrogen removal efficiency when the feed was high in sTKN and had a low sCOD:sTKN. Further, the impact of a seed inoculum on cell growth and protein synthesis can be explained by performance metrics such as yield and production rate (Table 2). Thus, a starting inoculum enriched from an existent community in the food-processing wastewater can provide microbial populations able to grow using the carbon sources and nutrients present, thus enhancing the rate of mixed-community SCP production.

### 3.5 Distinct microbial communities enriched from wastewater batches of variable quality

The most enriched SCP genera in reactors included *Azospirillum, Rhodobacter, Lactococcus*, and *Novosphingobium* (Figure 4). Except for *Lactococcus*, the other three main genera belonged to the phylum *Proteobacteria*, which is the predominant phylum reported in microbial community-based SCP production systems (Vethathirri et al., 2021). Although *Firmicutes* was the most prevalent phylum in the soybean processing wastewaters, *Proteobacteria* became dominant in the enriched microbial biomass in the reactors. Therefore, the massive immigration of bacteria present in the influent did not determine the structure of the microbial community in the reactors, contrary to what has been reported for full-scale activated sludge plants (Dottorini et al., 2021). A similar observation was made in a full-scale membrane bioreactor plant where the influent bacterial populations contributed little to relative abundances of the core activated sludge community (Zhang & Meng, 2021).

**Figure 4.**
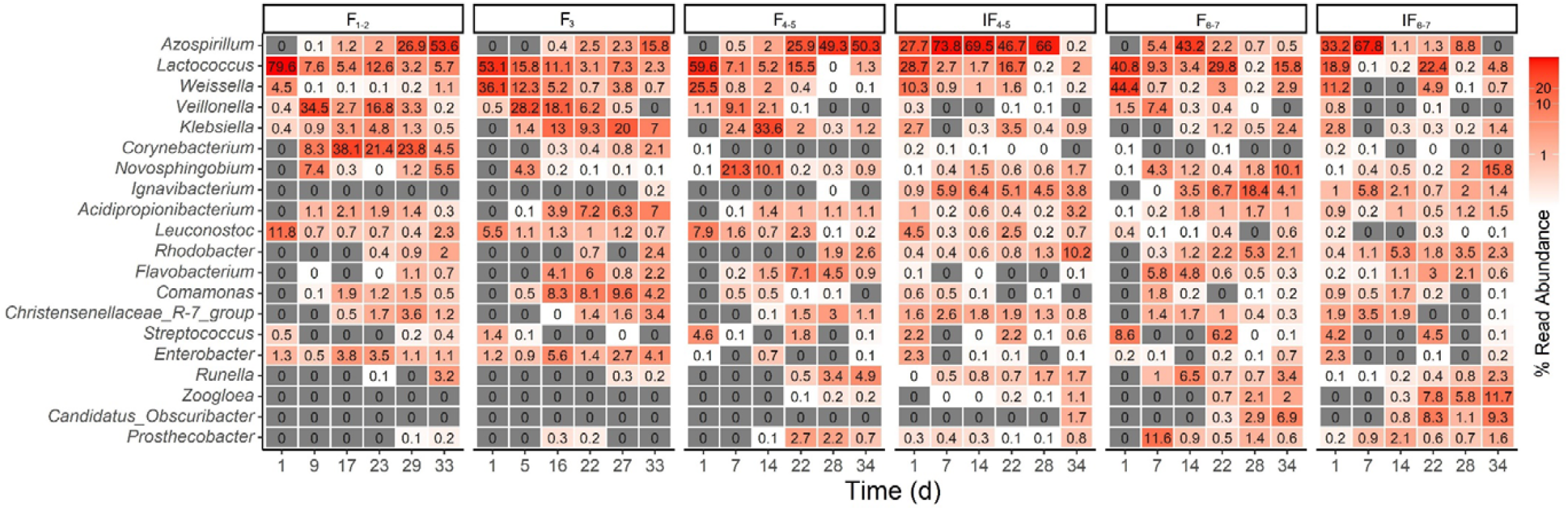
Microbial characterization based on 16S rRNA gene amplicon sequencing of community-based SCP biomass produced from soybean processing wastewaters. The temporal dynamics of the 20 most abundant bacterial genera across all reactors are shown. Four reactors (F_1-2_, F_3_, F_4-5_, and F_6-7_) were initiated without inoculum and two (IF_4-5_ and IF_6-7_) received a 1.5-L inoculum from reactor F_3_ (subscript numbers indicate the wastewater batch used as feed). Heat maps were generated using 16S rRNA gene amplicon v3-v4 data; ASVs were grouped at the required taxonomic level and ranked with the most abundant ASV on top. Additional reactor samples were analyzed for high temporal resolution (Figure S6-S8).

The observed shift in the microbial community composition in our study was likely due to selection conditions imposed by the available nutrients in the soybean-processing wastewaters and the operational conditions in the reactors (Santillan et al., 2020a; Santillan et al., 2019b; Seshan et al., 2021). Further, the reactors that operated without a starting inoculum (F_1-2_, F_3_, F_4-5_, F_6-7_), and received different batches of soybean processing wastewaters as feed, differed in the three most abundant bacterial genera after 34 days (Figure 4), despite having a similar initial community dominated by *Lactococcus* and *Weisella*. This highlights the effect of variable chemical characteristics of the influent wastewaters on the selection of core SCP communities in the bioreactors. Similarly, the two reactors that received the same sludge inoculum (IF_4-5_, IF_6-7_) with high relative abundances of the genera *Azospirillum* and *Lactococcus*, had enriched different bacterial genera after 34 days. These two reactors were fed different batches of soybean-processing wastewaters, which had a different chemical composition (Table 1). The two main genera in the reactors on d34 were *Rhodobacter* and *Ignavibacterium* and *Novosphingobium* and *Zoogloea*, respectively, also differing from the main genera in the influent wastewaters. Likewise, *Lactococcus*, which was the predominant genus in influent wastewaters, was not among the core SCP microbial communities except for two reactors (F_1-2_ and F_6-7_) that were fed with different wastewater batches. Similarly, *Weissella*, which was the second most abundant bacterial genus in the soybean wastewaters, was not enriched in any of the reactors. Taken together, these data suggest that organisms present in the influent wastewaters at low relative abundances (which may be below the limit of detection) can be enriched if provided appropriate selective conditions.

### 3.6 Enriched taxa with potential applications other than SCP production

Microbial community-based SCP production using food processing wastewaters may also lead to the enrichment of taxa that can produce valuable compounds or have further applications in addition to microbial protein production (Vethathirri et al., 2021). Following the shift from high (110 mg/L) to low (31 mg/L) sTKN in reactor F_1-2_, the relative abundance of *Azospirillum* increased from 2.0% to 53.6%. Their dominance could be due to low oxygen and nitrogen levels, which favor the growth of diazotrophs (Lee et al., 2015). FISH imaging further confirmed the presence of rod-shaped *Azospirillum* cells in the SCP community (Figure 5). Some taxa within the *Azospirillum* genus can both fix nitrogen (Fukami et al., 2018) and promote plant growth by synthesizing phytohormones and other valuable compounds that improve root growth and uptake of nutrients. A similar enrichment of *Azospirillum* to that of reactor F_1-2_ was observed in reactor F_4-5_, which was also started without inoculum and was fed with two wastewater batches of low sTKN content. *Azospirillum* was further identified as the most abundant bacterial genus in reactor F_3_, despite it having received wastewater with a high sTKN level of 110 mg/L throughout the 34-d experiment. This could have been due to the higher TSS level of 1160 mg/L from day 18 onwards in F_3_ compared to the other reactors, which translated into higher biomass production and use of available nitrogen. Reactor IF_4-5_ received a seed inoculum in the beginning and favored *Azospirillum* until d28, but its relative abundance decreased drastically from 66% to 0.2% after the change to a different wastewater batch of higher quality. The latter helped in the enrichment of the other two genera, *Rhodobacter* and *Acidipropionibacterium*, in reactor IF_4-5_ from d28 to d34. The different enrichment patterns of *Azospirillum* in the six reactors emphasize the impact of chemical variability of wastewaters from the same food processing industry on microbial composition of microbial community-derived SCP. Further, nitrogen-fixing bacteria like *Azospirillum* could help in the SCP production from nitrogen-limited wastewaters, with the added value of a low dissolved oxygen requirement. *Acidipropionibacterium* was among the three most enriched genera in reactors F_3_ and IF_4-5_. Previously named *Propionibacterium*, taxa within this genus have a wide range of applications in the food and pharmaceutical industries due to their ability to produce vitamin B12, trehalose, and bacteriocins via cell growth and synthesis of organic acids (Piwowarek et al., 2018). Some species within this genus, including *Acidipropionibacterium acidipropionici* (present in the reactors), play a major role in the food industry as either bio-preservatives or potential probiotics because of their generally recognized as safe (GRAS) status (Deptula et al., 2019). The genus *Novosphingobium* that was prevalent in reactor IF_6-7_ includes emerging diazotrophic strains capable of promoting rice cultivation in the absence of a nitrogen source (Rangjaroen et al., 2017). *Novosphingobium nitrogenifigens and Novosphingobium sediminicola*, which were both identified at the species level in the reactor, have been shown to accumulate polyhydroxyalkanoates when growing on nitrogen deficient pulp and paper-mill effluent (Addison et al., 2007), and to achieve high N levels facilitating the growth of the sugarcane plant in N-free sands (Muangthong et al., 2015), respectively. Another abundant organism from the same reactor, *Zooglea resiniphila*, has been found to enhance the removal of major resin acid toxicants in pulp and paper-mill effluents after bioaugmentation (Yu & Mohn, 2002). The most enriched species in reactor IF_4-5_, *Rhodobacter gluconicum*, was reported to be a dominant soil microbe involved in electricity generation in plant microbial fuel cells (Maddalwar et al., 2021). *Ignavibacterium*, another abundant genus specific to this reactor, was reported to be involved in the degradation of aromatic compounds in petrochemical wastewater effluents (Wang et al., 2021).

**Figure 5.**
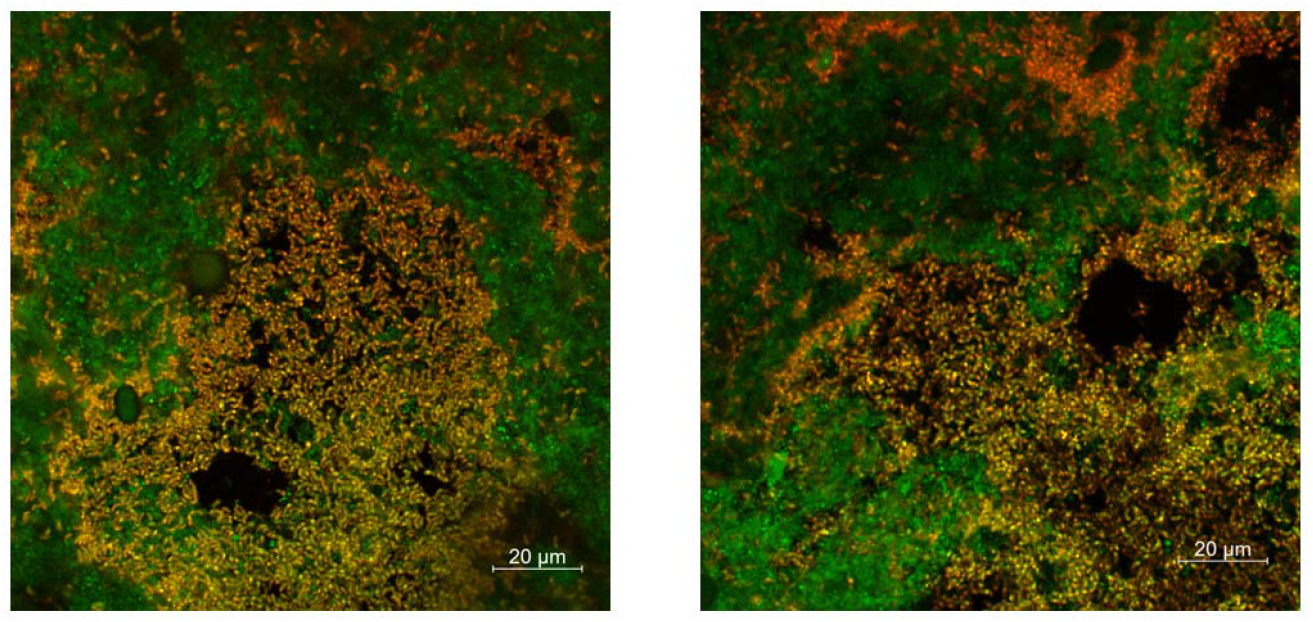
FISH images from reactor IF_4-5_ on d18 showing *Azospirillum* (orange) and all bacterial (green) cells. Cell abundances are consistent with 16S rRNA gene amplicon sequencing data for *Azospirillum* in IF_4-5_ on d18 indicating a relative abundance of 75.3%. Reactor IF_4-5_ had an inoculum from prior SCP production (reactor F_3_) and received batches 4 and 5 of soybean processing wastewater as feed. FISH images showing *Azospirillum* were also obtained from other reactors (Figure S9).

### 3.7 Limitations of the study

The overall protein production rates attained in this study were lower than those reported in prior studies on SCP production. The highest protein production rate occurred in reactor F_6-7_ (0.011 g SCP/L_R_/d) and was about 19 times lower than the rate reported for photosynthetic bacteria using artificial sugar wastewater with a COD:TN of 5 (Cao et al., 2021). Similarly, the highest biomass yield, also from reactor IF_6-7_ (0.089 g TSS/g sCOD_feed_), was ten times lower than that of photosynthetic nonsulfur bacteria produced using a volatile fatty acid-based medium with a COD:TN of 17 (Peng et al., 2022). These differences could be due to the use of synthetic wastewater with a controlled low C:N in those studies compared to the real food-processing wastewaters of varying C:N used here. Further, the sCOD:sTKN values of the soybean wastewaters batches used in this study (Table 2) were much higher than the recommended elemental C:N values (10 to 20) for microbial community-based SCP production (Vethathirri et al., 2021). The frequency of feeding was low, since time was required for the existent microbial communities in the wastewaters to grow before a new batch of feed was added. Additionally, a well-defined mixed-culture inoculum selected on specific substrates could have enhanced the production rate and yield. Hence it is plausible that a longer enrichment period on soybean-processing wastewaters would have resulted in a more effective inoculum.

### 3.8 Future directions to enhance SCP production from food-processing wastewaters

Further optimization of operational parameters to enhance SCP production and, specifically, accumulation of essential amino acids, is desirable. We observed an average of 92% of sCOD and 73% of sTN removal (Table 2). The current feeding scheme was chosen to provide adequate time for building microbial biomass from soybean processing wastewater in SBRs. Future experiments starting with a previously enriched sludge inoculum may employ a more frequent feeding regime to increase the supply of carbon and nitrogen for cell protein synthesis. A similar strategy increased the production rate of intracellular polyhydroxyalkanoates tenfold, using volatile fatty acids as feed under aerobic conditions (Valentino et al., 2014). Additionally, microbial communities can produce other valuable compounds like vitamins and co-enzyme Q10 (Peng et al., 2022) through bioconversion of nutrient-rich synthetic wastewaters. Therefore, process parameters may be tweaked to allow the production of value-added microbial products as well as single-cell protein.

## 4. Conclusions

- Soybean-processing wastewater can be converted into SCP meeting aquaculture feed requirements through a mixed-community bioreactor bioconversion approach.
- SCP can be produced using the microbial communities already present in the food-processing wastewaters, although SCP production is higher when reactors are inoculated with a suitable mixed-community culture.
- Variable sTKN levels and sCOD:sTKN in the food-processing wastewaters result in distinct SCP microbial communities, regardless of the similar microbial composition of the influent.
- Taxa at low abundance in the wastewaters can grow into core SCP producing organisms under suitable process conditions.
- Microbial community-based bioconversion of food-processing wastewaters into SCP can enrich for taxa with the potential to promote plant growth, have probiotic effects, and produce valuable compounds like vitamins.

## Data availability

DNA sequencing data are available at NCBI BioProjects PRJNA832086. See supplementary information for details about chemical analysis, microbial protein yield and production, nutrient removal efficiency and HRT estimation, chemical characteristics of bioreactor effluent and influent, rarefaction plots for 16S rRNA gene sequencing data, biomass protein content (as % dry weight), temporal amino acid profiles in reactors, amino acid profile in dry biomass of wastewater, and temporal dynamics of the 20 most abundant genera in each reactor.

## Author Contributions

RSV, ES and SW conceived the study. RSV and ES designed the experiment. SW obtained the funding for the study. RSV performed the experiment. RSV and SST did the laboratory chemical analyses. HYH and ES performed the molecular work. ES did the bioinformatics analyses. RSV and ES interpreted the data, generated the results, and elaborated the main arguments in the manuscript. RSV, ES and SW wrote the manuscript. All authors reviewed the manuscript.

## Acknowledgements

This research was supported by the Singapore National Research Foundation (NRF) and Ministry of Education under the Research Centre of Excellence Program, and the NRF Competitive Research Programme (NRF-CRP21-2018-0006) “Recovery and microbial synthesis of high-value aquaculture feed additives from food-processing wastewater”. DI Drautz-Moses provided support for the 16S rRNA gene amplicon library preparation and sequencing pipelines employed.

