## Supplementary Information for "Microbial community-based production of single cell protein from soybean-processing wastewater of variable chemical composition"

*Correspondence to:

Water quality parameters were measured in accordance with Standard Methods (APHA-AWWA-WEF, 2005) and targeted chemical oxygen demand (COD) (Standard Methods 5220 D) and nitrogen species (ammonium, nitrite, and nitrate ions) using spectrophotometric tests (Hach Company, Loveland, Colorado, USA) and Ion Chromatography (Standard Methods 4500-$NH_3$ for ammonium; 4110 B for nitrate and nitrite). Total organic carbon (TOC) and total Kjeldahl nitrogen (TKN) were also measured in influent samples using a TOC-L analyzer (Shimadzu Corp., Kyoto, Japan). Total suspended solids (TSS) and volatile suspended solids measurements were performed according to Standard Methods (APHA-AWWA-WEF, 2005). Prior to chemical analysis, effluent samples were filtered through a 0.2-µm pore size filter and the filtrate was stored at 4ºC for less than one week.

*Microbial protein production and yield, and nutrient removal efficiency*

The production rate (g product/L/d) and yield (g product/g substrate) parameters were calculated to evaluate the overall performance of reactors (Vethathirri et al., 2021). The biomass yield was calculated as the ratio of total suspended solids production and the total amount of sCOD or sTKN available through the feeding of wastewaters as given by Eq. (1) and Eq. (2), where $TSS_n$ and $TSS_0$ represent the final and initial mass of dry biomass, respectively. Similarly, the protein yield was calculated as the ratio of total protein production and the total amount of sCOD or sTKN available through the feeding of wastewaters as given by Eq. (3) & Eq. (4), where $SCP_n$ and $SCP_0$ represent the final and initial mass of biomass protein, respectively. The biomass production rate (Eq. (5)) was calculated as the ratio of total suspended solids production and the reactor working volume ($L_R$) over the entire bioconversion period (n). Likewise, the protein production rate (Eq. (6)) was calculated as the ratio of total protein production and the reactor working volume over the bioconversion period. Further, average removal efficiencies of nitrogen ($R_{sTN}$) and carbon ($R_{sCOD}$) present in the wastewaters were estimated as given by Eq. (7) and Eq. (8), respectively, where m represents the number of wastewater batches used for a reactor under the feeding scheme of this study, and $sCOD_{effluent,m}$ and $sTN_{effluent,m,}$ respectively, refer to the sCOD and sTN available in the reactor before the subsequent feeding when m$^{th}$ wastewater batch was used in the previous feeding.

$$Biomass\ yield\_sCOD = \frac{TSSn - TSS_0}{sCODfeed} \quad \text{[g biomass / g substrate fed]} \tag{1}$$

$$Biomass\ yield\_sTKN = \frac{TSSn - TSS_0}{sTKNfeed} \quad \text{[g biomass / g substrate fed]} \tag{2}$$

$$Protein\ yield\_sCOD = \frac{SCPn - SCP_0}{sCODfeed} \quad \text{[g protein / g substrate fed]} \tag{3}$$

$$Protein\ yield\_sTKN = \frac{SCPn - SCP_0}{sTKNfeed} \quad \text{[g protein / g substrate fed]} \tag{4}$$

$$Biomass\ production\ rate = \frac{TSSn - TSS_0}{LR.n} \quad \text{[g biomass / L reactor working volume /d]} \tag{5}$$

$$Protein\ production\ rate = \frac{SCPn - SCP_0}{LR.n} \quad \text{[g protein / L reactor working volume /d]} \tag{6}$$

$$RsCOD = \{\frac{sCODfeed{,}1 - sCODeffluent{,}1}{sCODfeed1} * 100\ + \cdots \frac{sCODfeed{,}m - sCODeffluent{,}m}{sCODfeedm} * 100\}/m \tag{7}$$

$$RsTN = \{\frac{sTNfeed{,}1 - sTNeffluent{,}1}{sTNfeed1} * 100\ + \cdots \frac{sTNfeed{,}m - sTNeffluent{,}m}{sTNfeedm} * 100\}/m \tag{8}$$

The average protein content in biomass (as a percentage of dry weight) of the various batches of soybean processing wastewaters used on day 1 across the six reactors was estimated at 37.7%, the average value measured on two samples of soybean processing wastewaters (Figure S4). Hence, the initial microbial protein (d1) for each reactor starting without inoculum was estimated as 37.7% of its initial biomass (TSS) value. For reactors $IF_{4\text{-}5}$ and $IF_{6\text{-}7}$, which had a starting inoculum obtained from $F_3$ on d43 of its operation, their initial microbial protein on d1 was estimated from $F_3$-d43 protein value (Figure S2) and their respective initial soybean wastewater batch, accounting for mixing ratios. Based on the available data from reactors $F_{1\text{-}2}$ and $F_3$, their SCP yield and production rate were calculated using values of d1 and d29, and from d1 and d32, respectively; while their biomass yield and production rate were calculated from d1 and d33 values. All parameters for the remaining four reactors were calculated using d1 and d34 values.

*Hydraulic retention time*

Following the study design, when the effluent sCOD of a reactor was less than 400 mg/L, the biomass was left to settle for 60 min, after which 1.85-L of the supernatant effluent was removed. The reactor was then filled with same volume of soybean wastewater as feed. Reactors experienced different growth and sCOD removal rates due to the variable nutrient quality of the different soybean wastewater batches used to feed them. Additionally, as these reactors were designed for SCP accumulation, no biomass was removed from them during the course of the study. Consequently, there were different feeding intervals across bioreactors and time during the bioconversion process. Hence, an average hydraulic retention time (HRT) was estimated using Eq. (9) (Tchobanoglous et al., 2013).

$$HRT = \left\{\frac{V}{Q1} + \cdots \frac{V}{Qm}\right\} / m \ \text{[d]} \qquad (9)$$

Where, V represents the working volume of the reactor (L), m is the number of feeding episodes through 34 d period excluding the first time, and Qm refers to the flow rate of influent wastewater during the $m^{th}$ feeding episode (L/d).

**Table S1.** Chemical characteristics of bioreactor effluent and influent.

| **Reactor** | **Day** | **sCOD_effluent**[a] [mg/L] | **sCOD_feed**[b] [mg/L] | **sTN_effluent**[a] [mg/L] | **sTN_feed**[b] [mg/L] |
|---|---|---|---|---|---|
| $F_{1-2}$ | 4 | 3517 | 9038 | 21 | 110 |
| | 10 | 78 | 9038 | 15 | 110 |
| | 16 | 0 | 9038 | 22 | 110 |
| | 20 | 291 | 9038 | 60 | 110 |
| | 24 | 406 | 9038 | 10 | 110 |
| | 29 | 688 | 2334 | 16 | 31 |
| | 33 | 321 | 2334 | 5 | 31 |
| $F_3$ | 13 | 246 | 8945 | 55 | 110 |
| | 17 | 427 | 8945 | 12 | 110 |
| | 22 | 996 | 8945 | 51 | 110 |
| | 25 | 1200 | 8945 | 24 | 110 |
| | 31 | 1733 | 8945 | 37 | 110 |
| $F_{4-5}$ | 24 | 195 | 4180 | 7 | 28 |
| | 28 | 360 | 4180 | 16 | 28 |
| | 30 | 290 | 4180 | 5 | 28 |
| | 32 | 103 | 3474 | 4 | 34 |
| | 34 | 85 | 3474 | 6 | 34 |
| $IF_{4-5}$[c] | 18 | 226 | 4180 | 11 | 28 |
| | 22 | 491 | 4180 | 11 | 28 |
| | 24 | 231 | 4180 | 3 | 28 |
| | 28 | 280 | 4180 | 8 | 28 |
| | 30 | 380 | 4180 | 9 | 28 |
| | 32 | 148 | 3474 | 7 | 34 |
| | 34 | 108 | 3474 | 4 | 34 |
| $F_{6-7}$ | 24 | 202 | 2501 | 12 | 15 |
| | 28 | 270 | 2501 | 13 | 15 |
| | 30 | 230 | 2501 | 4 | 15 |
| | 32 | 218 | 7663 | 5 | 80 |
| | 34 | 298 | 7663 | 5 | 80 |
| $IF_{6-7}$[c] | 24 | 135 | 2501 | 2 | 15 |
| | 28 | 200 | 2501 | 4 | 15 |
| | 30 | 300 | 2501 | 2 | 15 |
| | 32 | 270 | 7663 | 4 | 80 |
| | 34 | 210 | 7663 | 7 | 80 |

[a] Effluent discharged after 1 h of settling, before feeding phase.

[b] Influent wastewater used right after effluent discharge.

[c] Reactors $IF_{4-5}$ and $IF_{6-7}$ started with 1.5-L inoculum from reactor $F_3$, while reactors $F_{1-2}$, $F_3$, $F_{4-5}$ and $F_{6-7}$ started without inoculum (subscript numbers indicate the wastewater batch used as feed).

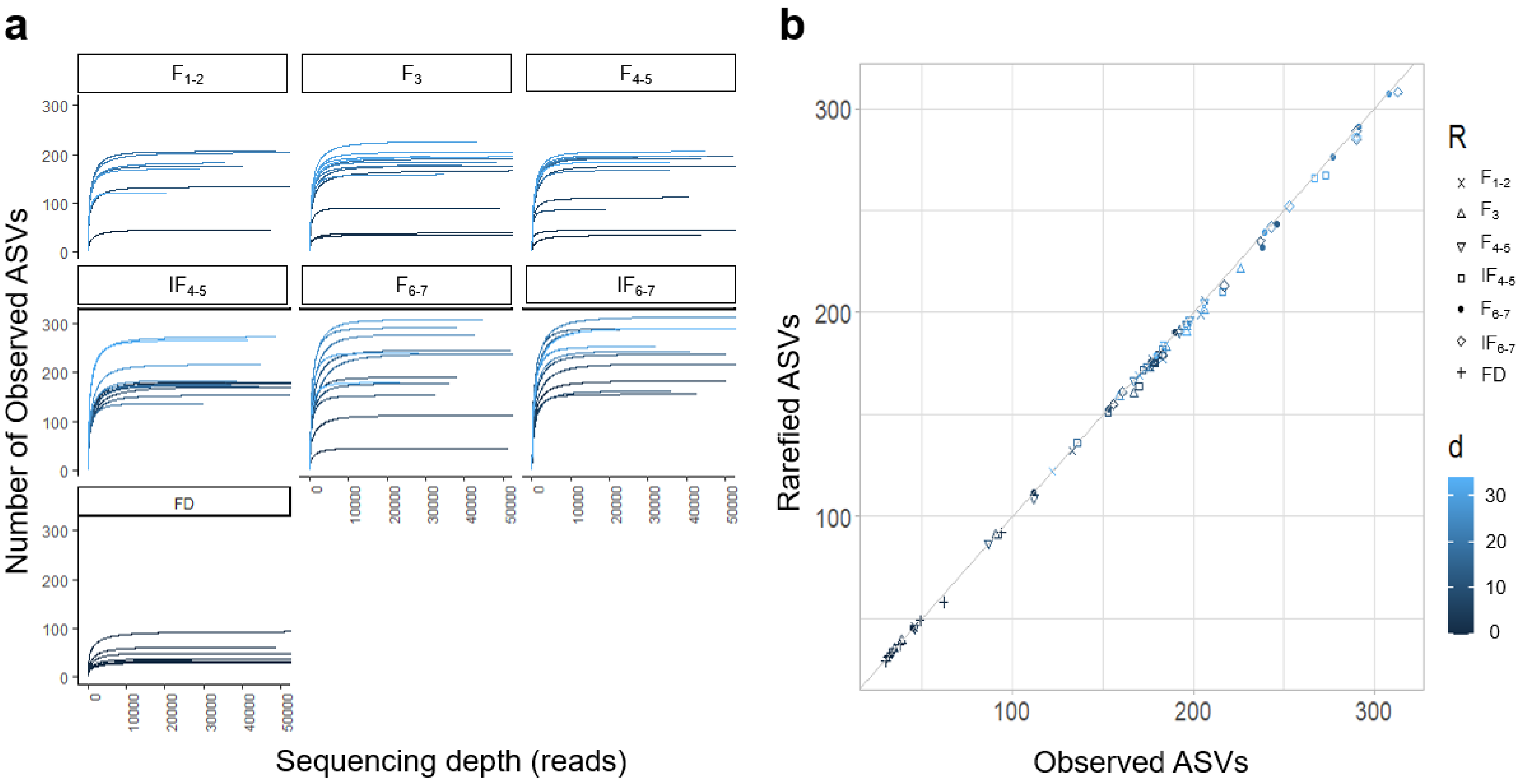


**Figure S1** Rarefaction plots for 16S rRNA gene sequencing data. (**a**) Rarefraction curves for reactors and soybean wastewater batches (FD) over time (**b**) Rarefied versus observed number of ASVs. Four reactors ($F_{1-2}$, $F_3$, $F_{4-5}$, and $F_{6-7}$) were initiated without inoculum and two ($IF_{4-5}$ and $IF_{6-7}$) received a 1.5-L inoculum from $F_3$. Subscript numbers indicate the wastewater batch used as feed.

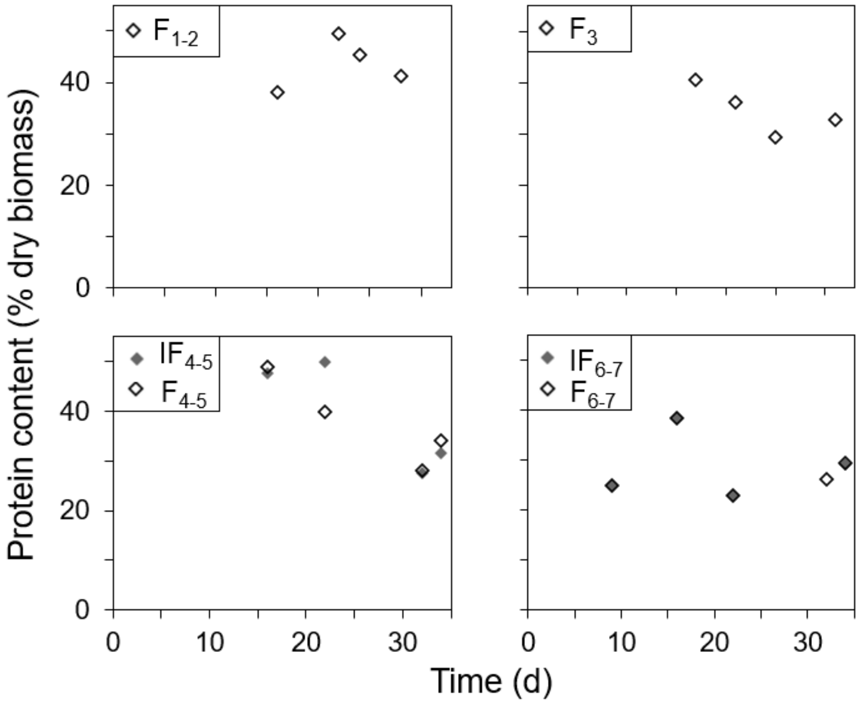


**Figure S2.** Biomass protein content (as % dry weight) in six SCP-enriched bioreactors determined using amino acid quantification via HPLC. Four reactors ($F_{1-2}$, $F_3$, $F_{4-5}$, and $F_{6-7}$) were initiated without inoculum and two ($IF_{4-5}$ and $IF_{6-7}$) received a 1.5-L inoculum from $F_3$. Subscript numbers indicate the wastewater batch used as feed.

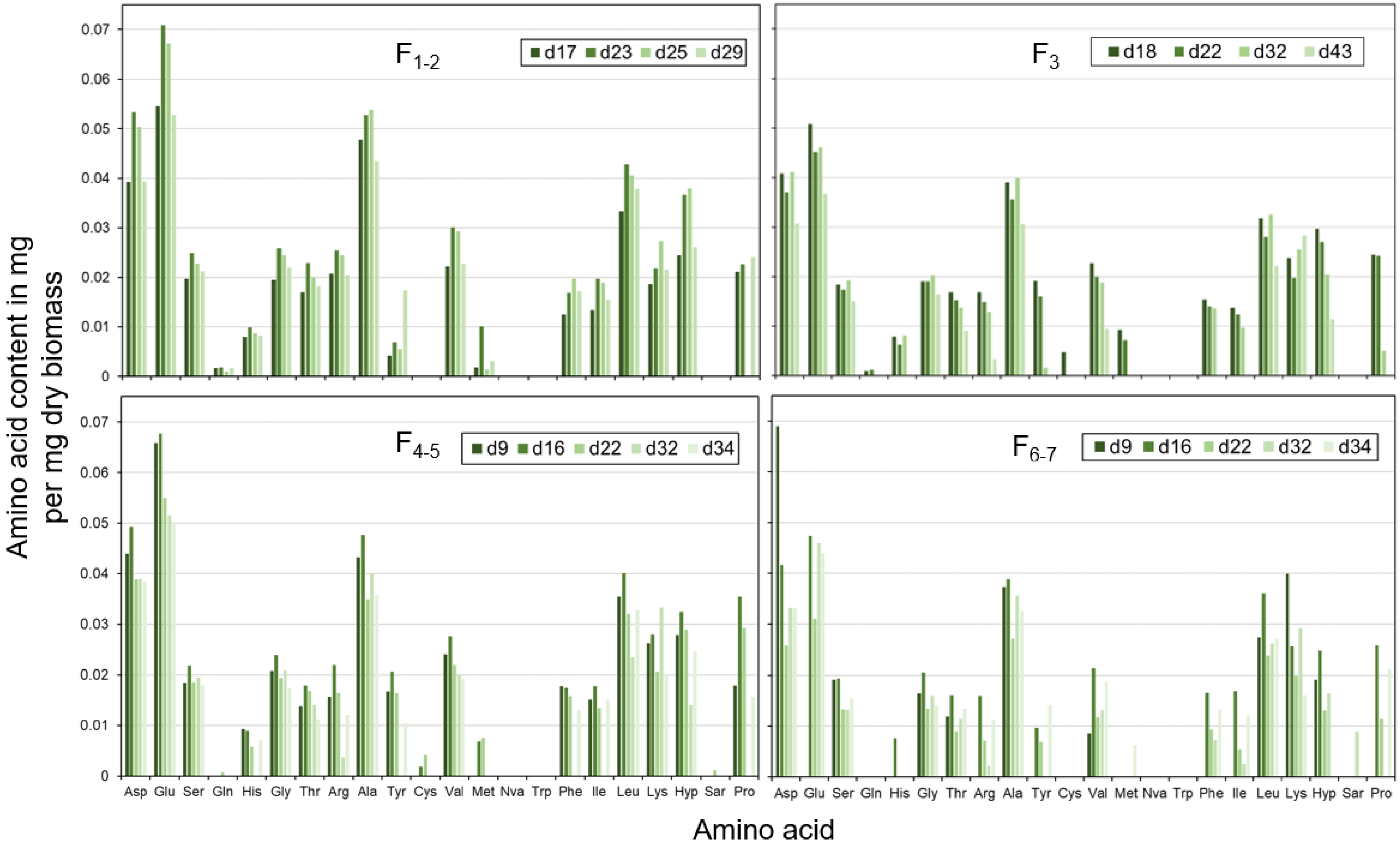


**Figure S3.** Temporal amino acids profiles (on days 9, 16, 22 and 34) in reactors operated without inoculum ($F_{1-2}$, $F_3$, $F_{4-5}$ and $F_{6-7}$). Subscript numbers indicate the wastewater batch used as feed.

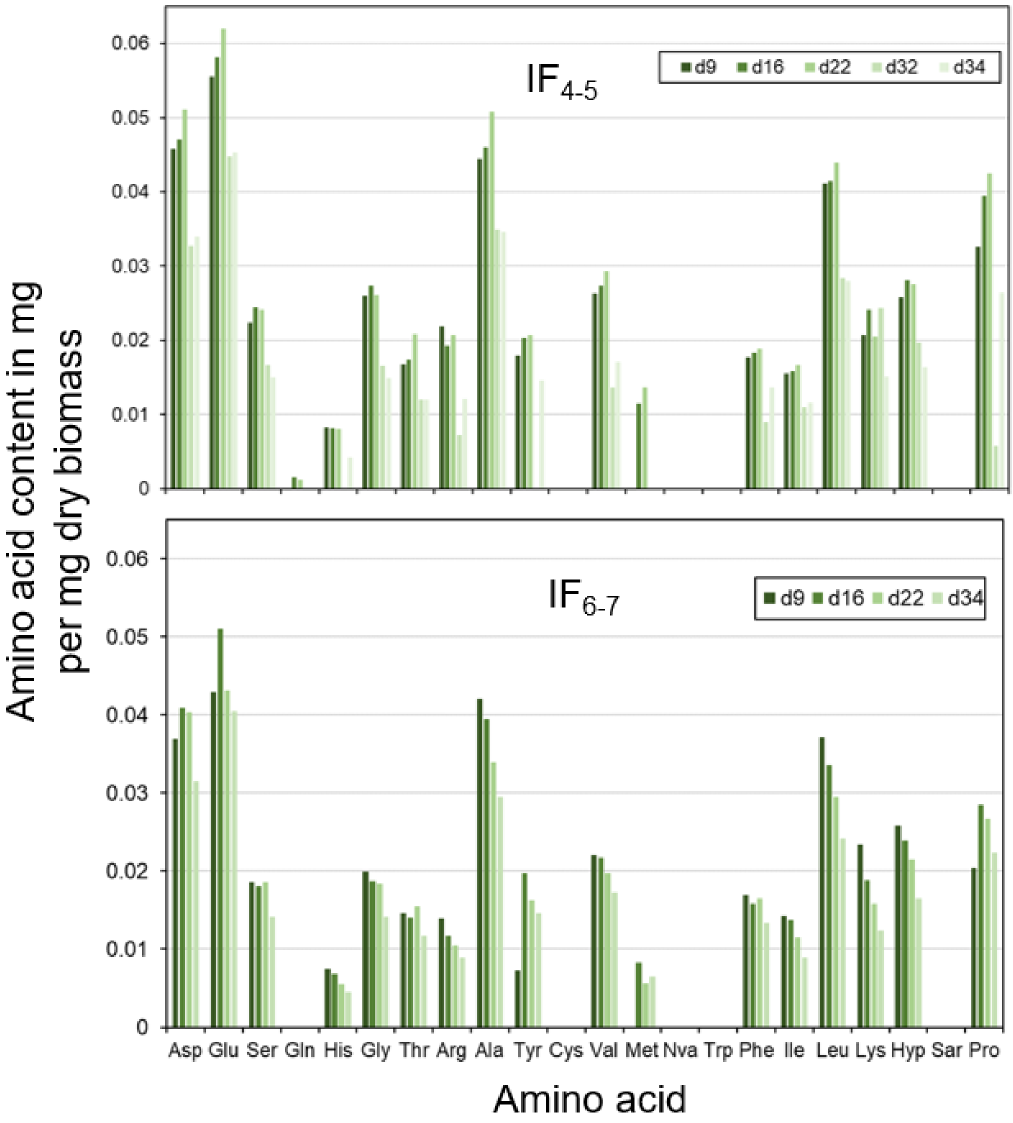


**Figure S4.** Temporal amino acids profiles (on days 9, 16, 22 and 34) in reactors that received an inoculum ($IF_{4\text{-}5}$ and $IF_{6\text{-}7}$). Subscript numbers indicate the wastewater batch used as feed.

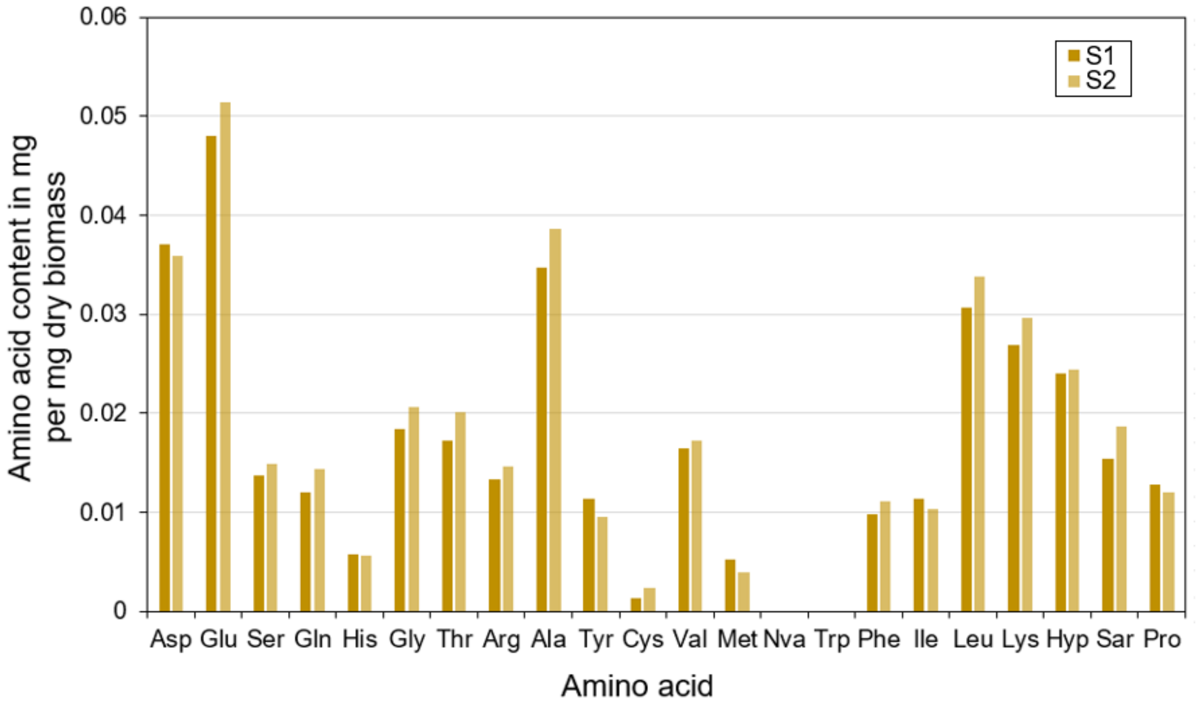


**Figure S5.** Amino acids in the dry biomass representing microbial communities in soybean processing wastewater used as feed for reactors. The average protein content in biomass (as a percentage of dry weight) of the various batches of soybean processing wastewaters used on day 1 for the six reactors was estimated using the average value measured in two wastewater samples (S1 and S2).

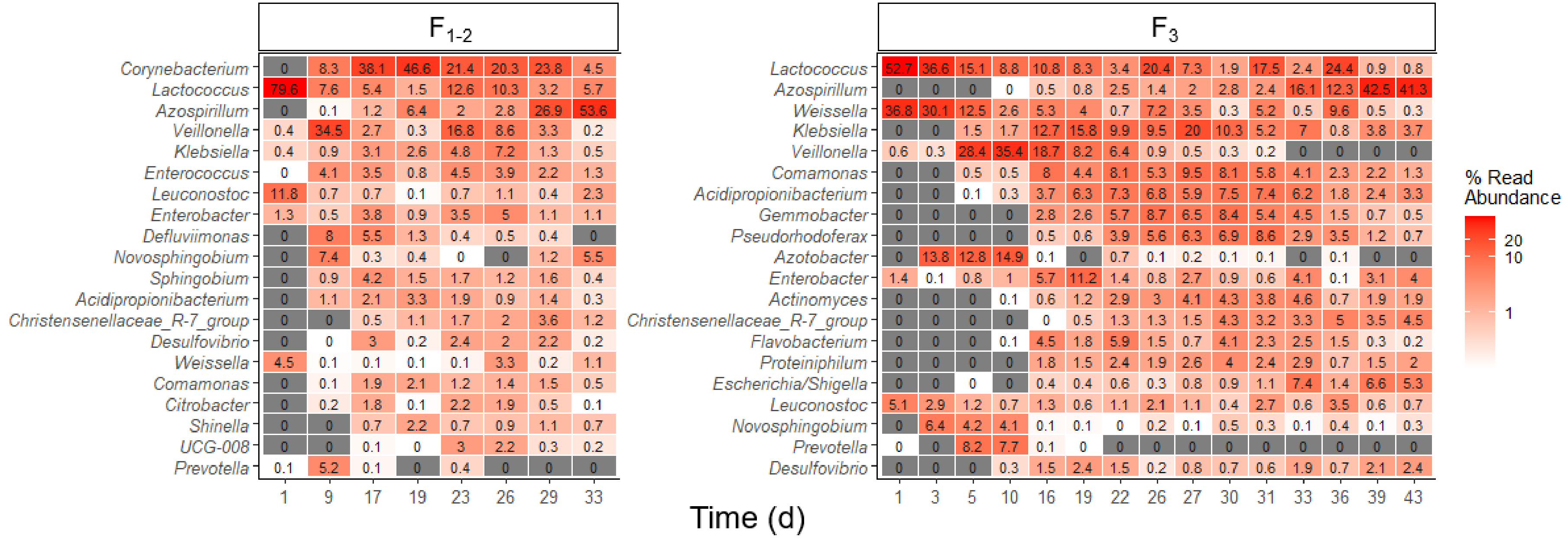


**Figure S6.** Temporal dynamics of the 20 most abundant genera in reactors $F_{1\text{-}2}$ and $F_3$, which started without inoculum and were operated for 34 d and 43 d, respectively. Reactors $IF_{4\text{-}5}$ and $IF_{6\text{-}7}$ had a 1.5-L starting inoculum taken from $F_3$ on d43. Heat maps were generated using 16S rRNA gene amplicon v3-v4 data, with the most abundant genera at the top. Subscript numbers indicate the wastewater batch used as feed.

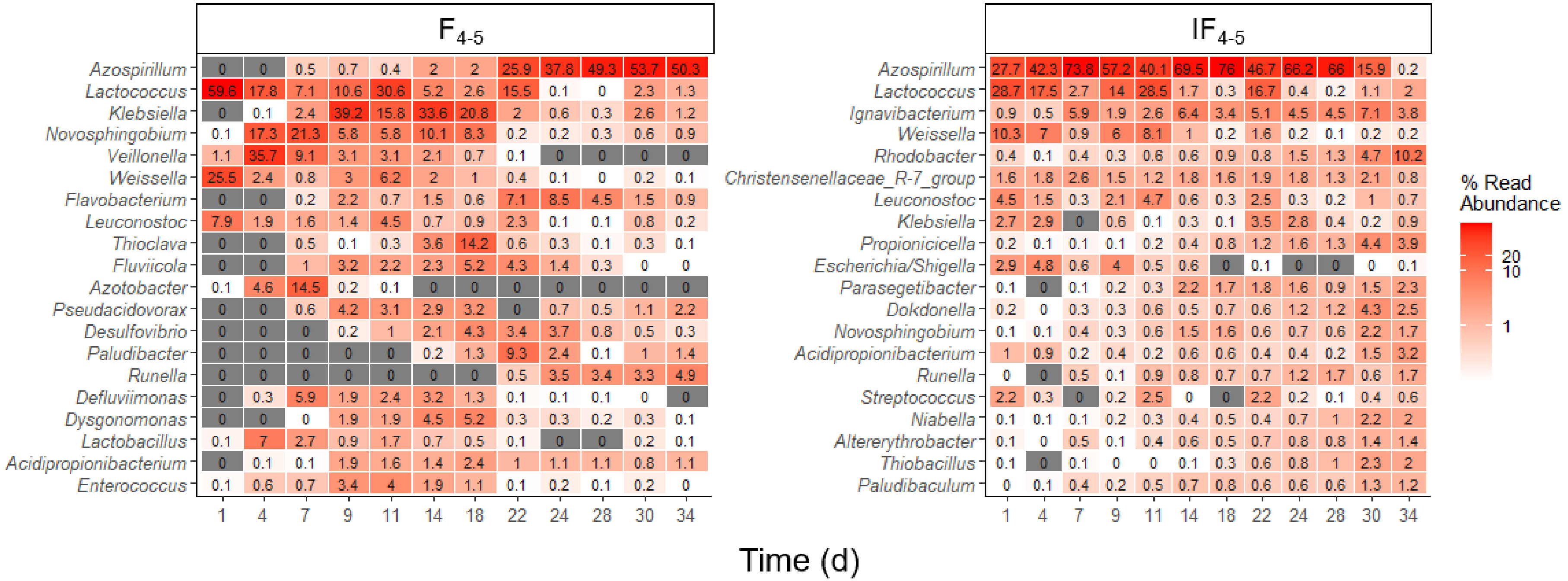


**Figure S7.** Temporal dynamics of the 20 most abundant genera in reactors $F_{4-5}$ and $IF_{4-5}$ operated for 34 d. Heat maps were generated using 16S rRNA gene amplicon v3-v4 data, with the most abundant genera at the top. Reactor $F_{4-5}$ was initiated without inoculum and reactor $IF_{4-5}$ received a 1.5-L inoculum from $F_3$ on d43. Subscript numbers indicate the wastewater batch used as feed.

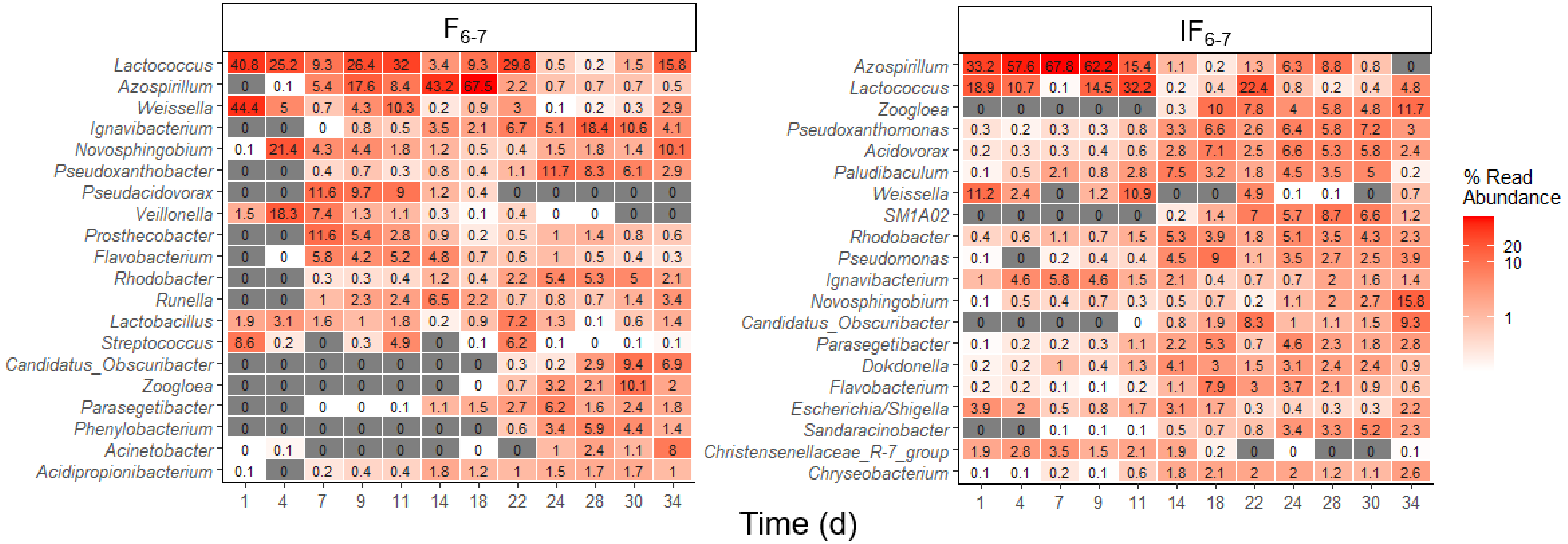


**Figure S8.** Temporal dynamics of the 20 most abundant genera in reactors $F_{6\text{-}7}$ and $IF_{6\text{-}7}$ operated for 34 d. Heat maps were generated using 16S rRNA gene amplicon v3-v4 data, with the most abundant genera at the top. Reactor $F_{6\text{-}7}$ was initiated without inoculum and reactor $IF_{6\text{-}7}$ received a using 1.5-L inoculum from $F_3$ on d43. Subscript numbers indicate the wastewater batch used as feed.

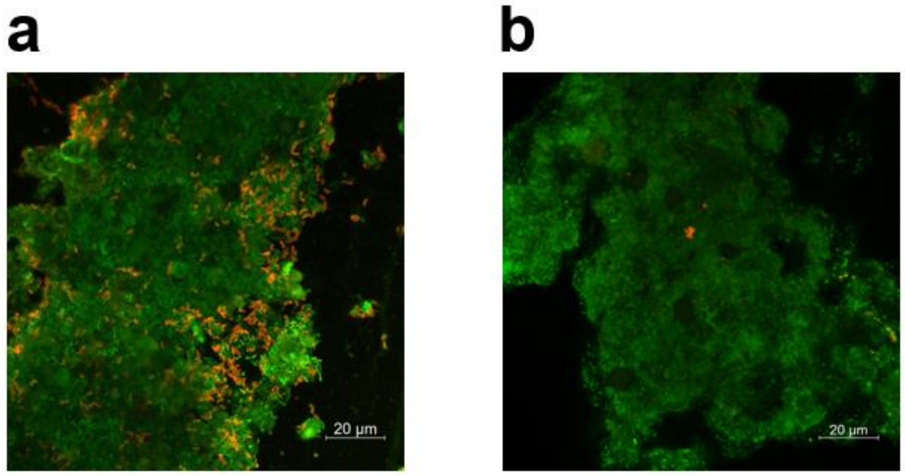


**Figure S9.** FISH images from **(a)** reactor IF$_{6-7}$ on d11 and **(b)** negative control sample showing *Azospirillum* (orange) and all bacterial (green) cells. Cell abundances are consistent with 16S rRNA gene amplicon sequencing data for for *Azospirillum* in IF$_{6-7}$ on d11 and a negative control sample, which indicated relative abundances of 15.2% and 0.3%, respectively**.** Reactor IF$_{6-7}$ had an inoculum from prior SCP production (reactor F$_3$) and received batches 6 and 7 of soybean processing wastewater as feed.
